# AMPK alters proteasome phosphorylation status and prevents persistent proteasome condensates

**DOI:** 10.1101/2025.07.01.662652

**Authors:** Jianhui Li, Conner Butcher, Kyle VanderVen, Mark Hochstrasser

**Author notes:** Corresponding author: Mark Hochstrasser. Co-corresponding author: Jianhui Li.

## Abstract

Proteasomes are large multiprotein complexes required for selective intracellular protein degradation, regulating numerous cellular processes and maintaining protein homeostasis and organismal health. In the budding yeast *Saccharomyces cerevisiae* grown under different glucose conditions, proteasomes undergo dynamic phase transitions between free and condensate states concomitant with nucleocytoplasmic translocation. Low glucose-induced cytoplasmic proteasome condensates are usually reversible but become persistent in the absence of AMP-activated protein kinase (AMPK). AMPK is important for proteasome condensate dissolution and proteasome nuclear reimport upon glucose refeeding of quiescent cells. Here we found that AMPK activities and the AMPK signaling pathway affect proteasome subunit phosphorylation, which correlates with the solubility and reversibility of proteasome condensates. Nuclear and cytoplasmic AMPK functions redundantly in proteasome condensate dissolution. AMPK interacts with the proteasome regulatory particle in an AMPK activity-independent manner. At least 50 kinases and phosphatases have been found to associate with the AMPK complex. Therefore, the prevention of persistent proteasome condensate formation by AMPK likely results from regulating the antagonistic effects of downstream kinases and phosphatases on proteasome phosphorylation. A mechanistic understanding of the downstream effector proteins of AMPK that directly regulate proteasome subunit phosphorylation will provide insights into how proteasome phosphorylation is linked to proteasome condensate regulation.

**Article summary:** Proteasomes undergo dynamic nucleocytoplasmic translocation and phase transitions in response to glucose starvation. AMP-activated protein kinase (AMPK) is important for cytoplasmic proteasome condensate dissolution and proteasome nuclear reentry in budding yeast cells upon glucose refeeding of quiescent cells. This study demonstrates that AMPK interacts with proteasomes, and the AMPK pathway regulates proteasome phosphorylation status and condensate solubility during reversible proteasome condensate formation. AMPK and the PP1 phosphatase dynamically regulate phosphorylation of multiple proteasome subunits. Therefore, the regulation of proteasome phosphorylation by AMPK is likely to be central to proteasome biomolecular condensate formation and dissolution.

## Introduction

The ubiquitin-proteasome system (UPS) is essential for selective protein degradation and regulates almost every aspect of cellular function, such as cell cycle progression and antigen processing (ROCK *et al*. 1994; TU *et al*. 2012). Dysfunction of the UPS is involved in the pathogenesis of many diseases, including many cancers, neurodegeneration, infection, inflammation, and developmental disorders (COUX *et al*. 2020). At the center of the UPS is the 26S proteasome, a large multiprotein complex comprising a proteolytic core particle (CP) and a regulatory particle (RP). The RP is assembled from lid and base subcomplexes (BARD *et al*. 2018).

Proteasomes undergo dynamic phase transitions and intracellular movement under various stress conditions (ENENKEL AND ERNST 2025). Stress can induce proteasomes to transition from a free state to a condensate-like state and can regulate proteasome nucleocytoplasmic translocation. The cellular regulation of proteasome phase transitions and intracellular movements is strongly conserved from yeast to plants and mammalian cells.

Depending on the specific stress conditions, reversible proteasome condensates have been observed in the nucleus of mammalian cells and the cytoplasm or perinuclear region of yeast cells (ENENKEL *et al*. 2022). In exponentially growing budding yeast *Saccharomyces cerevisiae*, proteasomes are highly enriched in the nucleus (PACK *et al*. 2014). When yeast enter stationary phase, a glucose starvation condition, most proteasomes are exported from the nucleus to form reversible proteasome condensates or proteasome storage granules (PSGs) in the cytoplasm (LAPORTE *et al*. 2008).

Upon glucose refeeding of such quiescent cells, proteasome condensates dissolve rapidly, and proteasomes are reimported into the nucleus within minutes (LAPORTE *et al*. 2008; BUTCHER *et al*. 2025). Cellular factors or conditions that modulate these movements and physical changes include ubiquitin (GU *et al*. 2017); cytosolic pH (PETERS *et al*. 2013); endosomal sorting complex required for transport (ESCRT) complexes (LI *et al*. 2019; LI AND HOCHSTRASSER 2022); membrane fusion protein Pep12 (VANDERVEN *et al*. 2025); proteasome shuttle factors Rad23, Dsk2, and Ddi1; and ubiquitin polymers (WAITE *et al*. 2024). By contrast, cellular factors required for proteasome condensate dissolution are poorly understood. Our previous studies demonstrated that AMP-activated protein kinase (AMPK) is required for proteasome condensate dissolution (LI *et al*. 2019). However, how AMPK regulates proteasome condensate dissolution is unclear.

AMPK is an evolutionarily conserved master regulator of cellular energy homeostasis in all examined eukaryotes (GHILLEBERT 2011). AMPK activation mediates a metabolic switch from an anabolic to catabolic state under stress conditions. Dysregulation of AMPK has been implicated in the pathogenesis of many human diseases, such as metabolic disorders, neurodegenerative diseases, and cancer (ASHRAF AND VAN NOSTRAND 2024). AMPK is a heterotrimeric complex, comprising a catalytic α subunit (Snf1 in yeast), a regulatory β subunit, (the three yeast paralogs–Gal83, Sip1, and Sip2–determine the subcellular location of the AMPK complexes), and a regulatory γ subunit (Snf4) (COCCETTI *et al*. 2018). In yeast cells, Snf1 is activated by upstream kinases Sak1, Elm1, and Tos3 through the phosphorylation of threonine-210 (T210), equivalent to T172 in mammalian cells (JEON 2016), in the Snf1 activation loop (SUTHERLAND *et al*. 2003). The Reg1-Glc7 protein phosphatase 1 (PP1) reverses AMPK activation through T210 dephosphorylation (MCCARTNEY AND SCHMIDT 2001). Snf1 can also be negatively regulated by autoinhibition, which occurs through interaction of the Snf1 kinase with its regulatory domains (JIANG AND CARLSON 1996). Many drugs and compounds have been developed and identified that alter AMPK activity (STEINBERG AND CARLING 2019). Understanding the regulation of proteasomes by AMPK may offer a means to repurpose these FDA-approved AMPK-related drugs for proteasome regulation and proteasome dysfunction-related disease treatment.

Proteasome condensates are one of the hundreds of biomolecular condensates that can form under various specific conditions (NARAYANASWAMY *et al*. 2009). Biomolecular condensates play crucial roles in fine-tuning cellular processes, such as gene expression and intracellular signal transduction (JEON *et al*. 2025). At the cellular level, it is evident that AMPK plays an important role in regulating biomolecular condensate formation and dissolution. For instance, under arsenite stress, the Glc7 catalytic subunit of yeast PP1 relocates from the nucleus and assembles into cytoplasmic condensates, resulting in translational inhibition. The regulatory subunit Reg1 of PP1 and its cellular target Snf1 are important for regulating reversible Glc7 cytoplasmic condensate formation (SCHNELL *et al*. 2021). Notably, the RNA-binding protein Smaug1 forms cytosolic condensates, thereby inhibiting mRNA translation; activation of AMPK induces dissolution of the Smaug1 condensates, releasing mRNA for translation, whereas inhibition of AMPK locks Smaug1 in the condensed state (THOMAS *et al*. 2025).

Our previous studies demonstrated that persistent proteasome condensates are formed in the absence of AMPK when budding yeast are grown under low glucose conditions (LI *et al*. 2019). On the other hand, the UPS can also regulate AMPK (ZUNGU *et al*. 2011). Here, we show that the AMPK signaling pathway regulates proteasome condensate (PSG) dissolution. AMPK complexes interact with proteasomes and maintain proteasome condensate solubility under low glucose conditions. Snf1 kinase activity is necessary for regulating proteasome condensate dissolution, which correlates with altered proteasome phosphorylation status. Our findings thus illuminate a cellular pathway of dynamic proteasome phase transitions regulated by AMPK.

## Results

### Proteasome condensates gradually lose reversibility without AMPK

Proteasome condensates (PSGs) are reversible (LAPORTE *et al*. 2008). They dissolve rapidly in a repetitive “contact and release” manner at the nuclear periphery, where proteasomes reimport to the nucleus within 15 minutes upon glucose recovery (BUTCHER *et al*. 2025). AMPK is required for proteasome condensate dissolution upon glucose refeeding after cells are starved for four days in low-glucose medium (LI *et al*. 2019). To understand the role of AMPK in regulating proteasome condensate dissolution, we examined and quantified the percentage of cells with proteasome condensates after one and four days in low glucose and after glucose refeeding following these starvation treatments. As expected, proteasome condensates were reversible in wild-type (WT) cells under low glucose conditions on both day one and day four (Figure 1). By contrast, in the AMPK deletion mutants *snf1Δ* and *snf4Δ*, proteasome condensates rapidly dissolved upon glucose addition on day one but became persistent by day four of starvation (Figure 1). Parallel results were observed using GFP-tagged CP (Pre10), RP base (Rpn2), and RP lid (Rpn5) subunits. These data indicate that AMPK is important for maintaining the reversibility of PSGs following prolonged glucose limitation.

**Figure 1.**
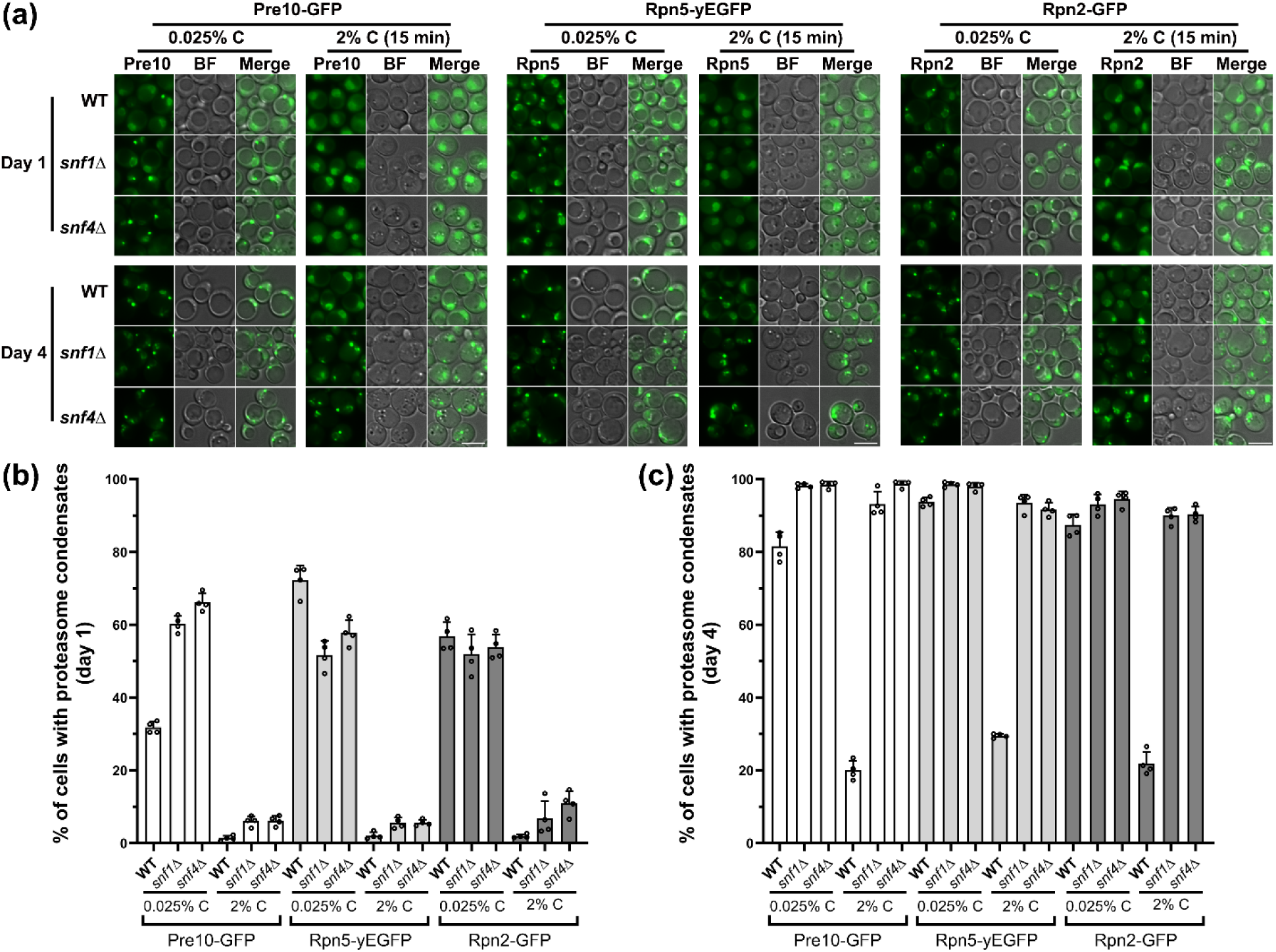
Proteasome condensates (proteasome storage granules or PSGs) gradually lose reversibility in the absence of AMPK under low glucose conditions. (a) Fluorescence microscopic images of the CP subunit Pre10-GFP, the lid subunit Rpn5-yEGFP, and the base subunit Rpn2-GFP in WT and AMPK deletion mutant *snf1Δ* and *snf4Δ* strains grown under low glucose conditions for one or four days and after 15 min of glucose recovery. There are only one to two PSGs per cell when starved. BF: bright field (phase). Scale bar: 5 µm. Persistent proteasome condensates were formed in *snf1Δ* and *snf4Δ* cells on day 4 under low glucose conditions. (b, c) Quantification of proteasome condensates in WT, *snf1Δ*, and *snf4Δ* cells following low glucose growth and after 15 min glucose recovery on day one (b) and day four (c). Bar graph results were plotted as mean±s.d. of the percentage of proteasome condensates in the indicated strain. Each sample has four data points that indicate the percentages. At least three repeats were conducted. More than 300 cells were counted for each sample.

### Persistent proteasome condensates are associated with abnormal proteasome phosphorylation status

To investigate how AMPK might control proteasome condensate reversibility, we used the Phos-tag gel mobility shift method to examine proteasome subunit phosphorylation status. Phos-tag molecules preferentially capture phosphomonoester dianions bound to serine, threonine, and tyrosine residues (KINOSHITA *et al*. 2006). Phosphorylation of proteasome subunits could be inferred by their slowed migration in Mn^2+^-Phos-tag SDS-PAGE gels. By anti-GFP immunoblotting of GFP-tagged subunits, we observed dynamic proteasome phosphorylation changes in cells under different glucose conditions; specifically, we switched the glucose concentration from 2% to 0.025% for one or four days and then switched the four-day-starved cells back to 2% glucose for 15 minutes to examine recovery (Figure 2).

**Figure 2.**
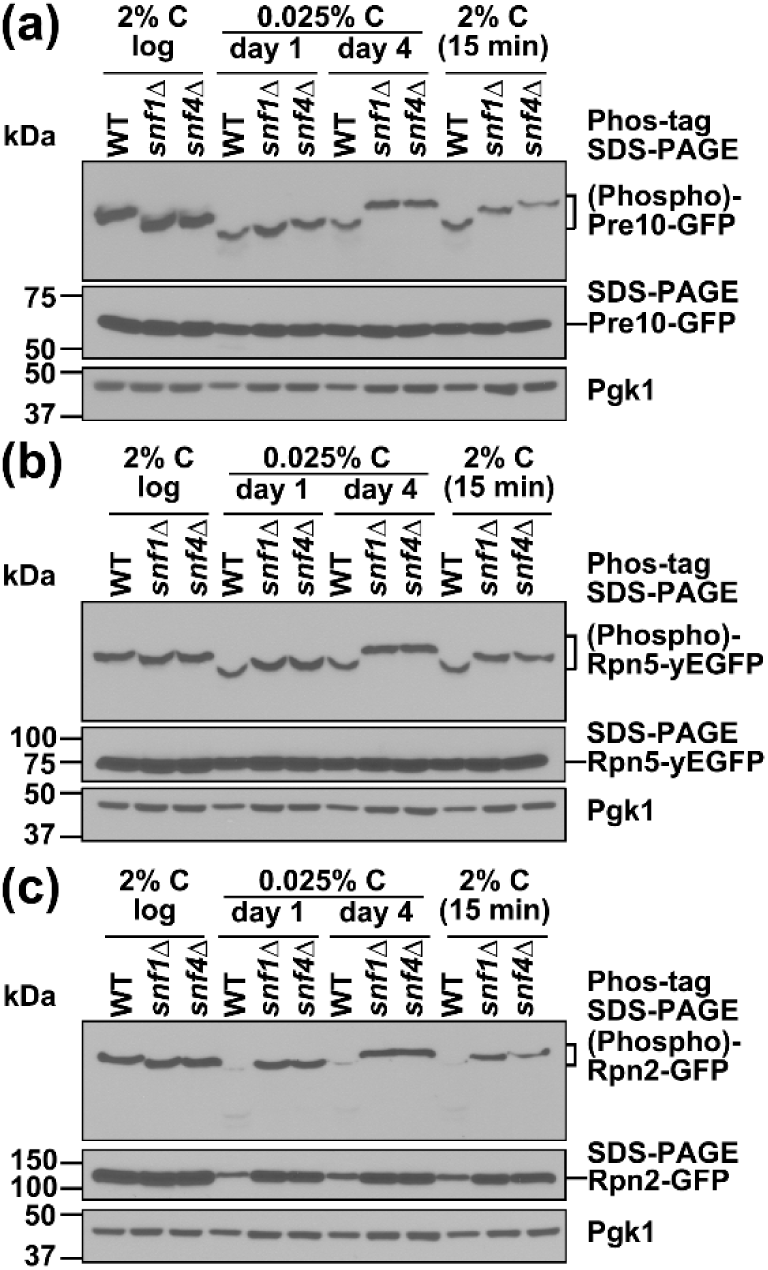
Highly phosphorylated proteasome subunits accumulate in AMPK deletion mutants after prolonged incubation in low glucose. Anti-GFP immunoblot analyses of proteasome subunit phosphorylation status (Phos-tag) and total proteasome subunit accumulation levels of Pre10-GFP (a), Rpn5-yEGFP (b), and Rpn2-GFP (c) in WT and AMPK mutant *snf1Δ* and *snf4Δ* cells. Phosphorylation retards the migration of proteins in the Phos-tag gels. Cells were cultured at the indicated glucose concentrations and harvested at the indicated times. Pgk1 (phosphoglycerate kinase-1) served as a loading control.

When isolated from exponentially growing (‘log’) WT cells in 2% glucose, all three tested GFP-tagged proteasome subunits migrated more slowly in Phos-tag gels than when grown under glucose-starved conditions. The CP subunit Pre10 is smaller than the lid subunit Rpn5 and the base subunit Rpn2, so phosphorylated Pre10 isoforms were better separated in the Phos-tag gels (Figure 2a). WT proteasome subunit phosphorylation decreased dramatically on day one of glucose limitation, based on the more rapid migration of the subunits in Phos-tag gels (Figure 2). While their degree of phosphorylation increased slightly on day four, it did not return to the levels seen in non-starved cells, and this also did not change detectably after 15 minutes of glucose refeeding. Therefore, proteasome phosphorylation undergoes striking changes in response to glucose availability. This correlates with proteasome nucleocytoplasmic translocation and proteasome transitions between free particles and proteasome condensates.

By contrast, proteasome subunits were underphosphorylated (faster migrating) in *snf1Δ* and *snf4Δ* cells growing exponentially in 2% glucose (Figure 2). Subunit phosphorylation decreased slightly by day one but increased by day four under low glucose to levels equivalent to WT levels or nearly so. These levels decreased slightly again during glucose recovery. Together, the results demonstrate that AMPK regulates proteasome phosphorylation status under changing glucose conditions. Moreover, the persistence of proteasome condensates observed in AMPK mutants correlates with abnormal proteasome subunit phosphorylation status.

### Snf1 kinase activity regulates proteasome condensate reversibility

Snf1 is activated by upstream kinases through phosphorylation of the conserved T210 residue in the Snf1 activation loop (SUTHERLAND *et al*. 2003). Lysine-84 (K84) and glycine-53 (G53) in Snf1 are important parts of the ATP-binding site and are essential for kinase activity (ESTRUCH *et al*. 1992). Mutations of T210 to alanine (snf1-T210A) or aspartate (snf1-T210D), or mutation of K84 to arginine (snf1-K84R) disrupt Snf1 kinase activity while G53 to arginine (snf1-G53R) mutation enhances Snf1 kinase activity (ESTRUCH *et al*. 1992). To test whether Snf1 activation through T210 and Snf1 kinase activity are essential for proteasome condensate dissolution and affect proteasome subunit phosphorylation status, we expressed WT *SNF1* or the aforementioned *snf1* mutants from the native promoter on a low-copy *CEN* plasmid in *snf1Δ* cells (Figure 3).

**Figure 3.**
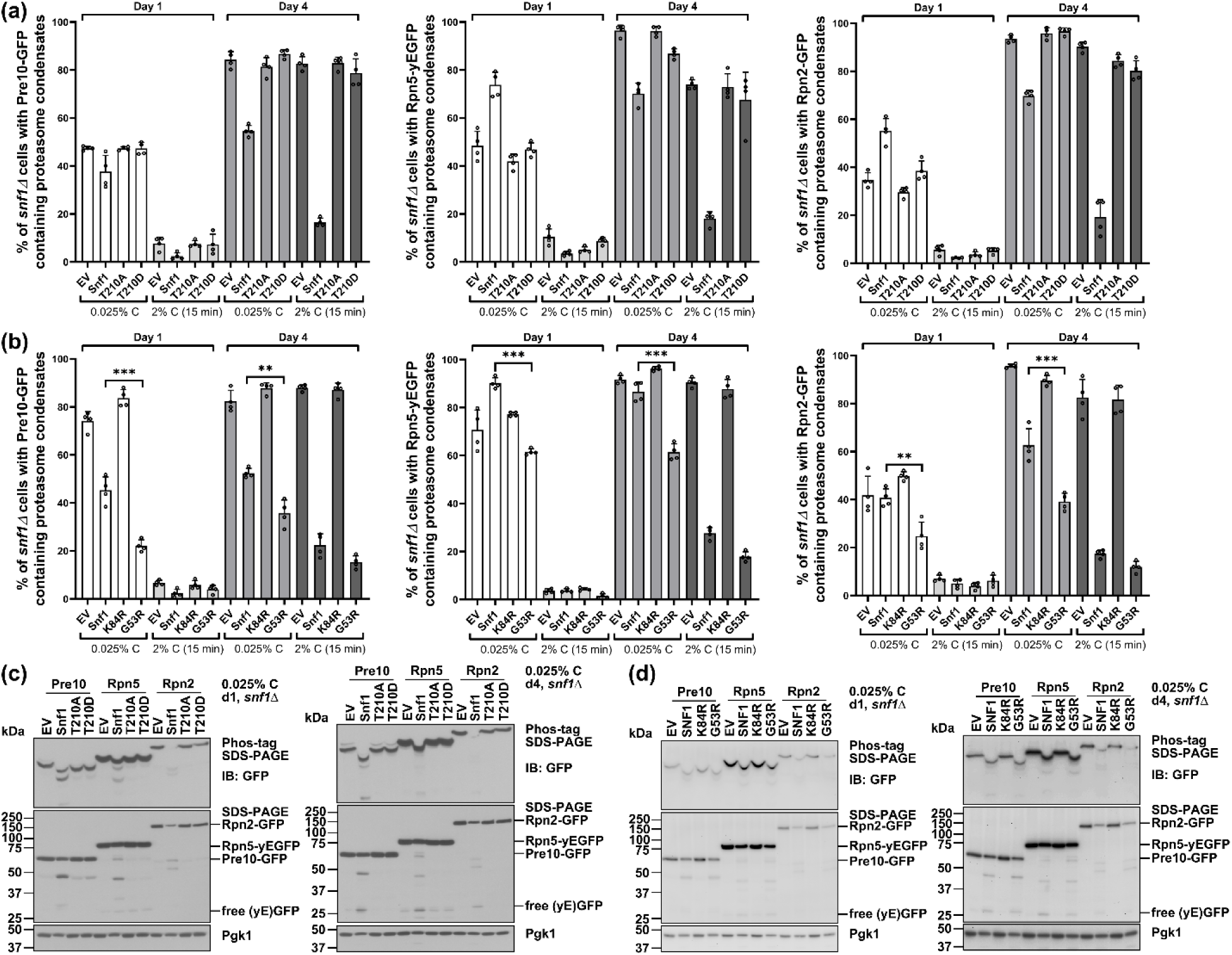
Snf1 activation and kinase activity are essential for both proteasome condensate dissolution and regulation of proteasome phosphorylation during glucose starvation and refeeding. (a) Percentage of cells bearing proteasome condensates tagged with Pre10-GFP, Rpn5-yEGFP, or Rpn2-GFP in *snf1Δ* cells carrying either an empty vector (EV) or plasmids with the indicated *snf1* alleles expressed from the native *SNF1* promoter. Mutants *snf1-T210A* and *snf1-T210D* are defective for kinase activity. (b) Percentage of cells bearing proteasome condensates tagged as indicated and grown under the indicated culture conditions. Mutant *snf1Δ* cells carried either an empty vector (EV) or plasmids with the indicated *snf1* alleles expressed from the native *SNF1* promoter. The snf1-K84R protein is kinase defective, while snf1-G53R has elevated kinase activity. \*\**P*<0.01 and \*\*\**P*<0.001 (one-way ANOVA analysis comparing snf1-G53R to Snf1 at indicated growth conditions. (c) Anti-GFP immunoblot analyses of proteasome subunit phosphorylation status (Phos-tag) and total proteasome subunit accumulation levels of Pre10-GFP, Rpn5-yEGFP, and Rpn2-GFP in cells from panel a. (d) Anti-GFP immunoblot analyses of proteasome subunit phosphorylation status and total proteasome subunit accumulation levels of Pre10-GFP, Rpn5-yEGFP, and Rpn2-GFP in cells from panel b. Cells were cultured under low glucose for about one day and four days, and subjected to glucose recovery for 15 min. Bar graph results were plotted as mean±s.d. of the percentage of proteasome condensates in the indicated strain. Each sample has four data points that indicate the percentages. At least three repeats were conducted. More than 300 cells were counted for each sample in panels a and b. Pgk1 served as a loading control in panels c and d.

Like *snf1Δ* cells with an empty vector (EV), abnormally persistent proteasome condensates were observed following glucose refeeding of cells that had been in low glucose for four days and expressed the inactive snf1-T210A, -T210D, or -K84R mutant proteins (Figure 3a and 3b). Correlating with the persistence of proteasome condensates, CP and RP proteasomal subunits remained more heavily phosphorylated in these mutants, comparable to *snf1* null cells (Figure 3c and 3d, Phos-tag SDS-PAGE gel blots). Microautophagy of proteasomes was also similarly impaired based on the reduced fragmentation or loss of these subunits compared to *snf1Δ* cells complemented with the WT *SNF1* plasmid. In contrast to the kinase-dead mutants, the percentage of *snf1-G53R* mutant cells (which have elevated kinase activity) bearing proteasome condensates was decreased relative to WT under low glucose conditions. Importantly, proteasome condensates were fully reversible in the *snf1-G53R* mutant upon glucose refeeding (Figure 3b), and proteasome subunit phosphorylation and autophagic degradation were unaffected (Figure 3d). These results indicate that Snf1 kinase activity regulates proteasome condensate reversibility, proteasome autophagy and subunit phosphorylation.

Proteasome condensate formation is a highly regulated cellular process that occurs alongside proteasome degradation by autophagy under low glucose conditions (LI *et al*. 2019). This helps ensure that normal proteasomes are sorted into reversible proteasome condensates while aberrant proteasomes are removed through autophagy (LI AND HOCHSTRASSER 2020). Snf1 has a dual role in regulating proteasome condensate dissolution and autophagic degradation (LI *et al*. 2019). Phosphorylation of T210 activates Snf1, whereas its dephosphorylation by PP1 inactivates the kinase. Reg1 is a regulatory subunit for the Glc7 phosphatase in the PP1 complex, and cells lacking Reg1 can no longer dephosphorylate T210 and exhibit constitutive phosphorylation of T210 and activation of Snf1 kinase activity (MCCARTNEY AND SCHMIDT 2001). Since elevated Snf1 kinase activity in the G53R mutant appeared to reduce the levels of proteasome condensates in low glucose and enhance their reversibility upon glucose refeeding (Figure 3b), we tested whether constitutively active Snf1 could also alter reversible proteasome condensates in *reg1Δ* cells. As predicted, a lower percentage of *reg1Δ* cells contained proteasome condensates compared to WT cells, and the condensates were smaller than in WT cells (Figure S1). To confirm that regulation of proteasome condensate formation by Reg1 is through the AMPK signaling pathway, we tested proteasome condensate reversibility in *snf1Δ reg1Δ* and *snf4Δ reg1Δ* cells. As expected, persistent proteasome condensates were formed in *snf1Δ reg1Δ* and *snf4Δ reg1Δ* cells (Figure S2). This demonstrates that *snf1Δ and snf4Δ* are epistatic to *reg1Δ* in regulating proteasome condensate dissolution.

These results indicate that constitutive Snf1 kinase activity caused by PP1 phosphatase inactivation promotes dissolution of proteasome condensates and that Snf1 activity levels can fine-tune proteasome condensate formation and dissolution.

### The regulatory β subunits Gal83 and Sip2 function redundantly in regulating proteasome condensate dissolution

The regulatory β subunits Sip1, Sip2, and Gal83 determine the subcellular activity of Snf1 in the vacuole, cytoplasm, and nucleus, respectively (JIANG AND CARLSON 1997; VINCENT *et al*. 2001). To understand which β subunit(s) is required for regulating proteasome condensate dissolution and which subcellular population(s) of AMPK regulates proteasome condensate dissolution, we examined proteasome condensates in single and double deletion mutants of the three β subunits. Proteasome condensates were reversible in all three single and two of the three double deletion mutants (Figure S3); the lone exception was the *sip2Δ gal83Δ* double mutant (Figure 4). Sip2 and Gal83 mediate cytoplasmic and nuclear Snf1 activity, respectively, suggesting that nuclear and cytoplasmic Snf1 activity are each sufficient to mediate proteasome condensate dissolution and nuclear reimport. The Sip1-Snf1-Snf4 complex is the only remaining Snf1 kinase complex in *sip2Δ gal83Δ* cells. However, previous work revealed low *in vitro* kinase activity due to the reduced levels of Sip1 in *sip2Δ gal83Δ* cells (NATH *et al*. 2002). Therefore, we cannot fully exclude the possibility that the persistent proteasome condensates in *sip2Δ gal83Δ* cells are due to overall reduced Snf1 kinase activity rather than loss of Snf1 activity at a particular subcellular site.

**Figure 4.**
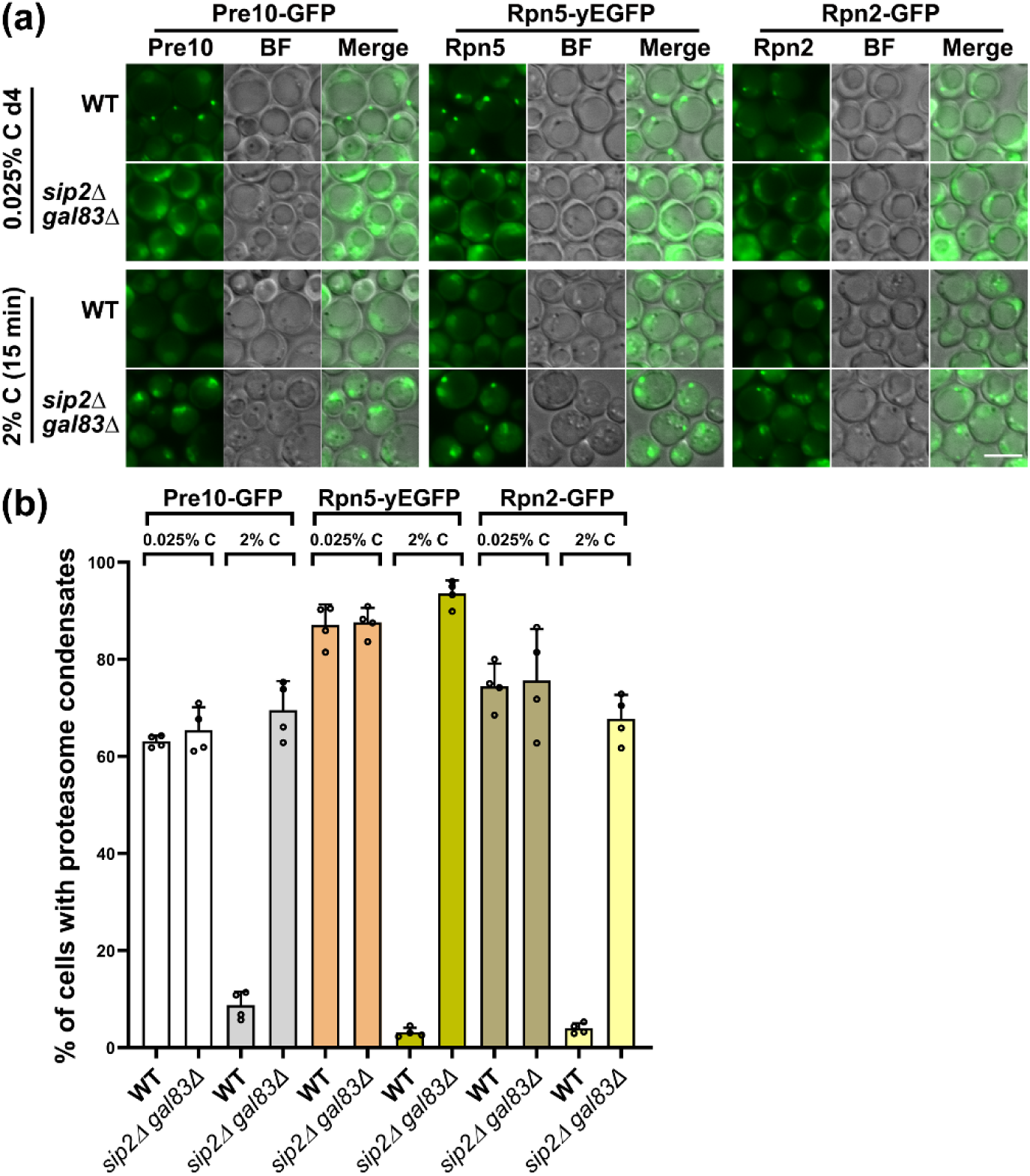
Nuclear and cytoplasmic AMPK complexes function redundantly in regulating proteasome condensate dissolution. (a) Fluorescence microscopy images of Pre10-GFP, Rpn5-yEGFP, and Rpn2-GFP in WT and *sip2Δ gal83Δ* cells grown under low glucose conditions for about four days and following glucose recovery for 15 min. BF: bright field. Scale bar: 5 µm. (b) Quantification of cells with proteasome condensates in WT and *sip2Δ gal83Δ* cells. The cells were from panel a. Bar graph results were plotted as mean±s.d. of the percentage of proteasome condensates in the indicated strain. Each sample has four data points that indicate the percentages. At least three repeats were conducted. More than 300 cells were counted for each sample.

### AMPK interacts with proteasome RP in a Snf1 kinase activity-independent manner

To examine whether AMPK interacts with proteasomes, we performed GFP-Trap coimmunoprecipitation in *snf1Δ* or *snf4Δ* cells expressing the GFP-tagged proteasome CP subunit Pre10 or RP subunit Rpn5 and plasmid-based expression of HA-tagged Snf1 or Snf4. Although *SNF1* and *SNF4* were both expressed from the *GPD* (*TDH3*) promoter, the levels of Snf4 protein appeared to be much higher than that of Snf1, possibly due to downregulation of Snf1 under low glucose conditions (Figure 5a, total cell lysates). Snf4 interacted with both the CP (Pre10-GFP) and the RP (Rpn5-GFP), whereas Snf1 only interacted detectably with the RP (Figure 5a, long exposure blot). WT Snf1, kinase activity-defective mutants T210A, T210D, and K84R, and elevated kinase-activity mutant G53R interacted with Rpn5 in a Snf1 kinase activity-independent manner (Figure 5b). Snf1 protein accumulation and that of the snf1 mutants varied and were negatively correlated with the kinase activity; specifically, Snf1 with the kinase-inhibiting T210A, T210D, or K84R mutations accumulated to higher levels than WT Snf1, while the hyperactive snf1-G53R kinase was at lower levels compared to WT Snf1 in total cell lysates (Figure 5b). No AMPK components have been detected in a previous tandem mass spectrometry analysis of WT strain without GFP tag (LI AND HOCHSTRASSER 2022). These results suggest that Snf1 kinase activity impacts Snf1 protein levels through a post-transcriptional mechanism but does not modulate Snf1-proteasome (RP) interactions.

**Figure 5.**
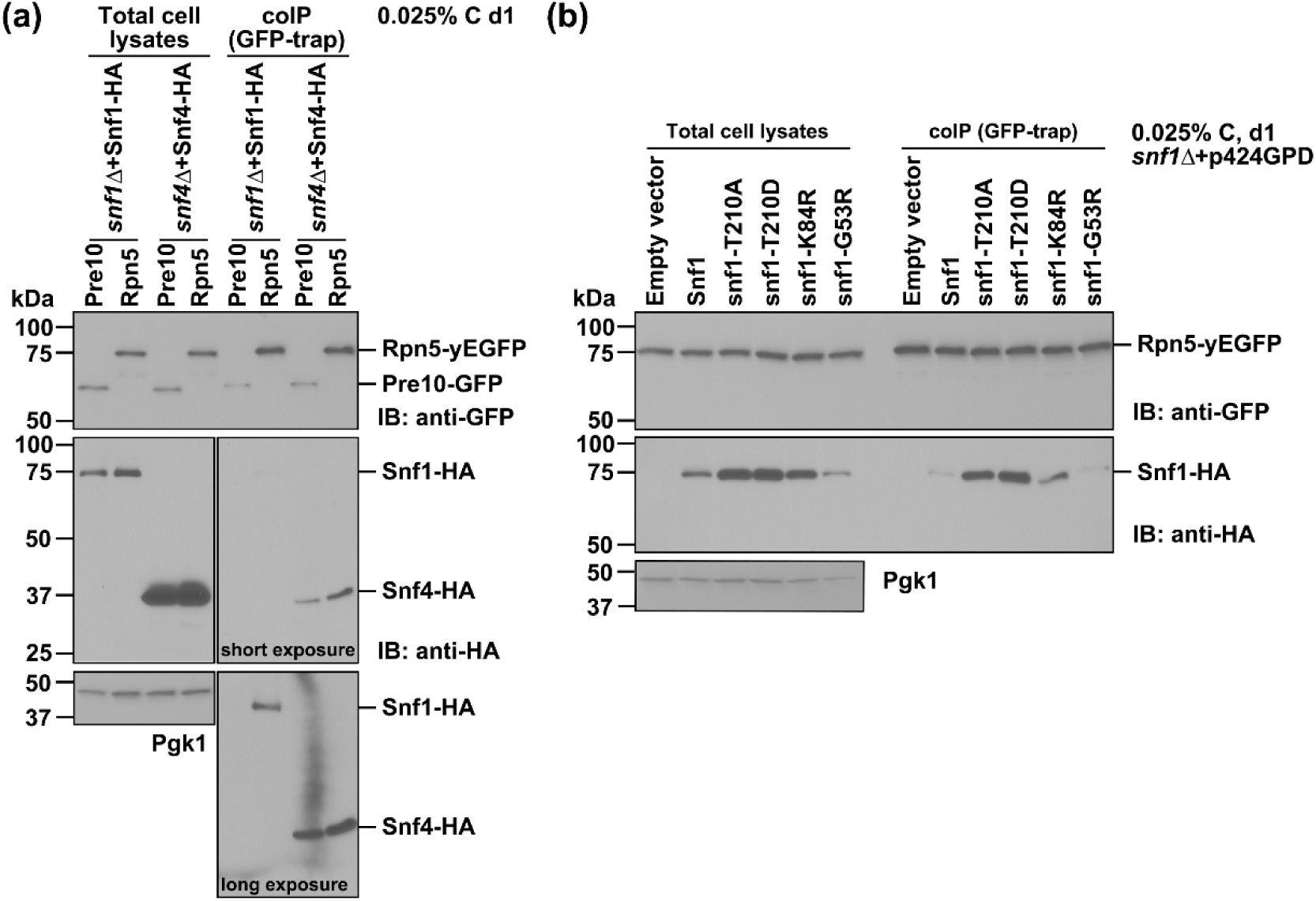
AMPK interacts with proteasomes in a Snf1 activity-independent manner. (a) Co-IP analyses of Snf1-HA and Snf4-HA with proteasomes in *snf1Δ* and *snf4Δ* cells carrying plasmids p424GPD-SNF1-HA and p426GPD-SNF4-HA, respectively. Snf4 interacted with both the CP subunit Pre10 and the RP subunit Rpn5, while Snf1 interacted detectably only with Rpn5, as shown in the long exposure blot. (b) Co-IP analyses of Snf1-HA and Snf1 kinase mutants with Rpn5. Cells were grown under low glucose conditions for about one day. Pgk1 served as a loading control. Relative to the amount that was loaded in immunoprecipitated samples, ∼0.83% of total cell lysates were loaded for the anti-HA and 6.67% for the anti-GFP blots.

We next purified Snf1 complexes from yeast cells expressing a 3xFlag-tagged Snf1 from the native chromosomal locus. By anti-Flag affinity purification followed by tandem mass spectrometry analysis, we could determine the Snf1 complex interactome under different growth conditions. Consistent with the co-immunoprecipitation results in Figure 5, Snf1 complexes interacted with the proteasomal CP, RP, and proteasome assembly chaperones in exponentially growing (log) cells, but only the RP complex copurified under low glucose conditions (Figure S4; Table S5). The lack of CP in the anti-Flag isolates after glucose limitation may reflect separation of CP and RP subcomplexes under these conditions (LI AND HOCHSTRASSER 2022) and a direct interaction of Snf1 only with the RP. Additionally, 37 kinases and 13 phosphatases were detected in the Snf1 interactome (Table S1). The results confirm the interactions between the Snf1 complex and the proteasomal RP and suggest the downstream effectors of AMPK signaling pathway regulate proteasome condensate reversibility.

### Persistent proteasome condensates in AMPK mutants behave as solid-like condensates

In the absence of AMPK in long-term glucose starved cells, proteasome condensates became resistant to dissolution following glucose refeeding and proteasomes were abnormally phosphorylated (Figure 2). More generally, protein phosphorylation plays important roles in regulating protein-protein interactions and can alter biomolecular condensate solubility and reversibility (SRIDHARAN *et al*. 2022; RANGANATHAN *et al*. 2023). Liquid-like, but not solid-like, biomolecular condensates can be dissolved by the solvent 1,6-hexanediol (1,6-HD), which interferes with their weak hydrophobic interactions. Therefore,1,6-HD is a useful tool to differentiate between liquid-like and solid-like condensates in cells (KROSCHWALD *et al*. 2017).

Using 1,6-HD treatments, we tested the physical properties of proteasome condensates (PSGs) in WT and AMPK mutant cells. Proteasome condensates in WT cells were readily dissolved by 1,6-HD after one day of glucose limitation (Figures 6a and 6b) but became less soluble after longer-term glucose starvation stress (Figures 6a and 6c). By contrast, proteasome condensates in AMPK deletion mutants were much more resistant to dissolving in 1,6-HD at both early and later times under glucose limitation, suggesting they had solid-like condensate properties (Figure 6). These data suggest that proteasome condensates lose solubility and behave more like solids or gels in cells lacking AMPK. Hence, an increase in solid-like proteasome condensates under low glucose stress correlates closely with loss of condensate reversibility and abnormal proteasome phosphorylation status in cells without AMPK (Figures 2 and 6).

**Figure 6.**
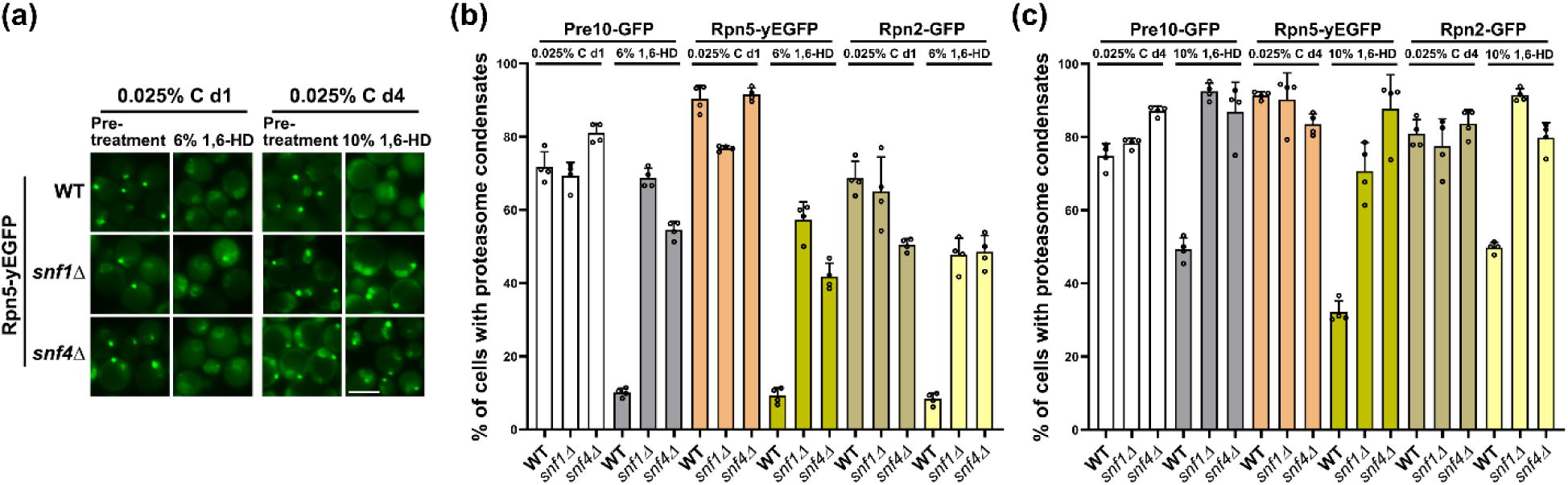
Solid-like proteasome condensates accumulate in mutant cells lacking AMPK. (a) Fluorescence microscopy images of Rpn5-yEGFP in WT, *snf1Δ, snf4Δ* cells grown under low glucose conditions for about four days and following 1,6-HD treatment for 20 min. Scale bar: 5 µm. (b) Quantification of proteasome condensates in WT, *snf1Δ*, and *snf4Δ* cells cultured under low glucose conditions for about one day followed by incubation of cells in 6% 1,6-HD for 20 minutes at room temperature (RT). (c) Quantification of proteasome condensates in WT, *snf1Δ*, and *snf4Δ* cells cultured under low glucose conditions for about four days followed by 10% 1,6-HD treatment for 20 minutes at RT. Bar graph results were plotted as mean±s.d. of the percentage of proteasome condensates in the indicated strain. Each sample has four data points that indicate the percentages. At least three repeats were conducted. More than 300 cells were counted for each sample. Proteasome condensate solubility in 1,6-HD was reduced over time under low glucose conditions.

In an attempt to identify the phosphorylation sites of proteasome subunits that lead to persistent proteasome condensates in cells, we performed gel phospho-staining and phospho-enrichment mass spectrometry of affinity-purified proteasomes from WT, *snf1Δ*, and *snf4Δ* cells. The Pro-Q® Diamond phosphoprotein gel stain allows direct, in-gel detection of phosphate groups attached to tyrosine, serine, or threonine residues. The phosphorylation status of several proteasome subunits, such as Rpt1, Rpn6, Rpn11, and Rpn12, was strongly impacted, with most showing decreased phosphorylation in *snf1Δ* and *snf4Δ* cells relative to WT cells, especially on day four under low glucose conditions (Figure 7). By contrast, proteasome subunit phosphorylation status was much less affected in *reg1Δ* cells, which have constitutively high AMPK activity (Figure S5). Therefore, loss of AMPK alters the phosphorylation status of multiple specific proteasome subunits in response to glucose starvation.

**Figure 7.**
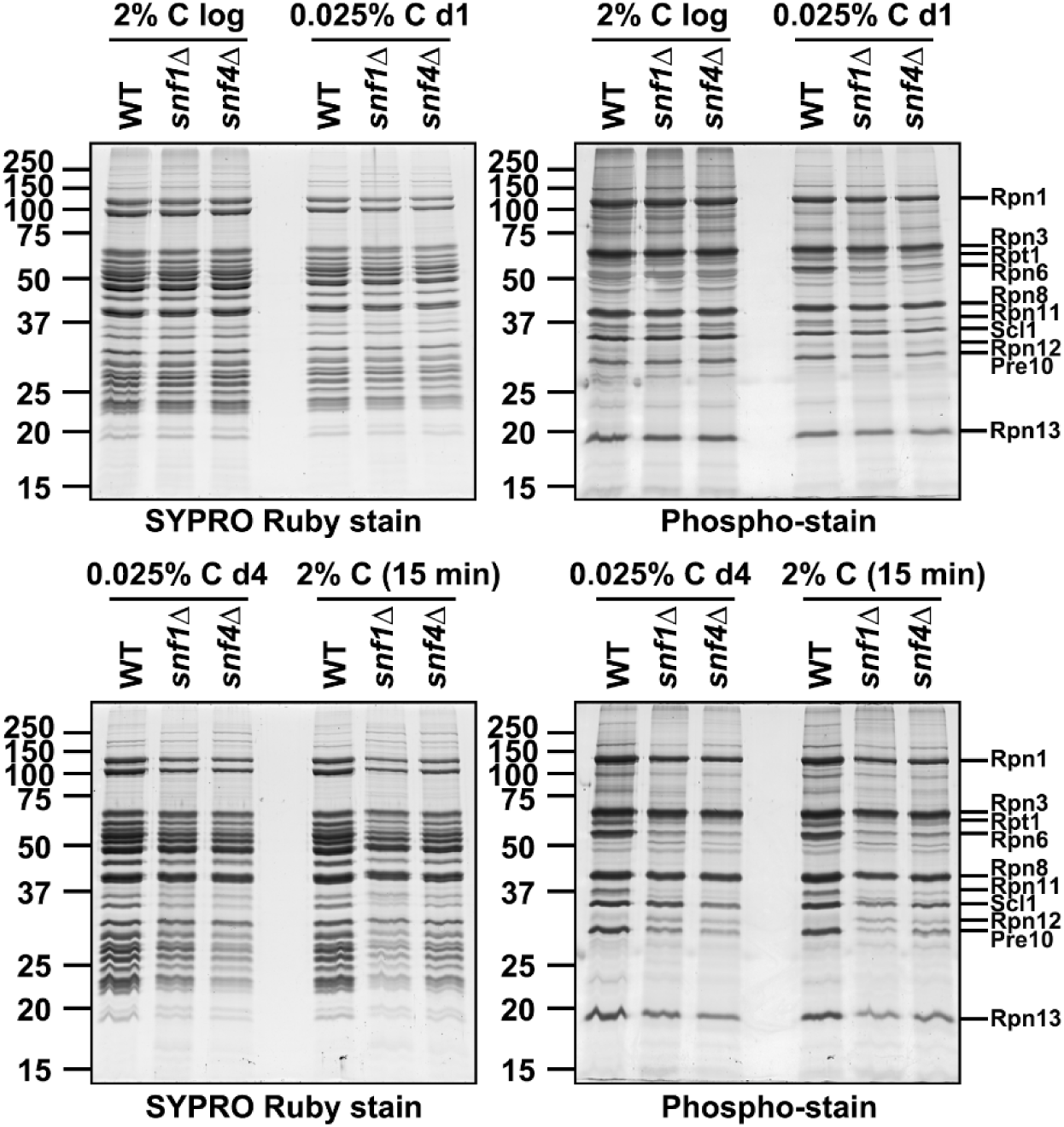
AMPK affects the phosphorylation of several proteasome subunits. Proteasomes were affinity purified (with an anti-Flag resin) from WT, *snf1Δ*, and *snf4Δ* cells expressing Rpn11-3xFlag under the indicated glucose growth conditions. Ten µg of proteasomes were resolved in 12% SDS-PAGE gels, stained with protein phospho-stain (right panels) followed by SYPRO Ruby stain (left panels). Pronounced changes in the phosphorylation status of several proteasome subunits were observed in cells without AMPK on day four under low glucose conditions. Proteasome subunits were labeled based on their molecular weights, so some labels might be inaccurate.

Phospho-enrichment mass spectrometry of purified proteasomes also revealed changes in phosphorylation status of individual proteasome subunits in AMPK null cells relative to WT when cells were starved for glucose (Figure S6). In most cases, the changes were in overall agreement with in-gel phospho-staining, such as for Rpn6 and Rpn11, but some discrepancies were seen (e.g., Pre10 phosphorylation was apparently lower in *snf1Δ* cells based on gel staining but was slightly increased based on the proteomic analysis). We investigated the impact on proteasome condensate dissolution of specific proteasome phosphorylation sites of several subunits (Rpn8, Rpn11, Rpn13, and Pre10) by mutating the relevant residues to alanine but failed to identify specific phosphorylation sites that were necessary. Given that proteasomes have 31 subunits that have been reported to be phosphorylated and hundreds of potential phosphorylation sites that may function redundantly (HIRANO *et al*. 2015), it is perhaps not surprising that we could not identify specific sites that recapitulate the phenotype of cells lacking AMPK.

## Discussion

In this study, we have demonstrated that Snf1 kinase activity is important for normal proteasome subunit phosphorylation patterns under low glucose growth conditions and for the solubility and reversibility of proteasome condensates (Figures 3, 6, and 7). The nuclear and cytoplasmic forms of Snf1 complexes together appear to be necessary for proteasome condensate dissolution, and either by itself is sufficient; the vacuole-localized Snf1 complex is neither necessary nor sufficient for condensate regulation (Figures 1 and 4). We observed a physical interaction between Snf1 and the proteasomal RP by co-immunoprecipitation and mass spectrometry; this interaction appears to be independent of Snf1 kinase activity (Figures 5 and S4).

To date, we have not been able to isolate Snf1 mutants that disrupt Snf1-RP interaction under low glucose conditions but retain AMPK activity. Therefore, it remains unclear how AMPK-proteasome interactions regulate proteasome trafficking and condensate properties. Snf1 complexes are unlikely to play a structural role in PSGs since glucose starvation-induced condensates are mainly composed of proteasomes and monoubiquitin (GU *et al*. 2017). Moreover, we did not detect direct binding between purified Snf1 complexes and proteasomes *in vitro* (not shown), indicating that additional factors may be involved in the physical interactions between these complexes in cells.

The physical interactions between Snf1 complexes and the proteasomal RP might suggest that AMPK can directly affect proteasome subunit phosphorylation. In our phosphorylation enrichment mass spectrometry analysis of purified proteasomes from WT and *snf1Δ* cells, we observed most proteasome subunits with reduced phosphorylation in *snf1Δ* cells on day four under low glucose (Figure S6). This is generally consistent with our phospho-staining results of purified proteasomes (Figure 7). Day four was also when persistent proteasome condensates were observed in cells lacking AMPK activity (Figures 1 and 3). Therefore, it is possible that specific AMPK-dependent proteasome subunit phosphorylations help maintain cytoplasmic proteasome condensates in a state capable of rapidly dissipating and allowing reimport of proteasomes back into the nucleus.

On the other hand, there were no clear differences in the phosphorylation patterns of purified proteasomes between WT and *snf1Δ* cells during exponential growth in high glucose or after one day in low-glucose conditions (Figure 7). These data using isolated proteasomes suggest it is less likely that AMPK acts directly on proteasomes since we should have detected more rapid responses to AMPK, and it should not take days to see changes in proteasome phosphorylation status. However, we did observe clear differences in proteasome phosphorylation profiles under changing glucose conditions when whole cell lysates were examined using Phos-tag gels and immunoblotting, and these changes correlated closely with loss of solubility of PSGs in 1,6-HD (Figures 2 and 6). As noted, the difference in samples of purified proteasomes versus total cell lysates and methodologies of phospho-stain versus Phos-tag gels may lead to different results (Figures 2 and 7).

Although we do not know the reason for the discrepancy between analyses of purified proteasomes analyzed by phospho-staining versus analysis of total cell lysates using Phos-tag gels, both methodologies suggest that AMPK functions indirectly on proteasomes by regulating other kinases and phosphatases that more directly impact the overall proteasome phosphorylation profile under the indicated growth conditions. This is consistent with the large number of phosphorylation sites found in proteasomes (HIRANO *et al*. 2015), and our detection of 37 kinases and 13 phosphatases in the Snf1 complex interactome that could function as part of the AMPK signaling pathway acting on proteasomes (Table S1). For example, Cdc14 phosphatase and Torc1 kinase have extensive network interactions with other kinases and phosphatases (BREITKREUTZ *et al*. 2010), and similar antagonistic effects of kinases and phosphatases on the overall proteasome phosphorylation status are also possible.

Phosphorylation plays a critical role in regulating formation and dissolution of biomolecular condensates, which are involved in diverse cellular and biological processes (SRIDHARAN *et al*. 2022). Phosphorylation alters protein conformations and interactions, which subsequently modulate condensate properties and biological functions (NOSELLA AND FORMAN-KAY 2021; RANGANATHAN *et al*. 2023). For instance, the N-terminal intrinsically disordered region of adenovirus 52K protein is phosphorylated at specific Ser residues, promoting liquid-like properties of viral 52K protein condensates and viral particle production (GRAMS *et al*. 2024). Previous studies have demonstrated that phosphorylation can alter proteasome function and activities (WANI *et al*. 2016; GUO *et al*. 2017; KORS *et al*. 2019; LIU *et al*. 2020). Our data demonstrate that phosphorylation is tightly linked to proteasome condensate solubility and reversibility (Figures 1, 2, 6, and 7). This is expected to alter proteasome functions, for example, by altering nuclear re-import dynamics of proteasomes upon glucose refeeding.

To identify the phosphorylation sites in proteasome subunits that are important for proteasome condensate dissolution, we performed TiO_2_ phospho-enrichment mass spectrometry analysis of purified proteasomes from WT and *snf1Δ* cells (Figure S6). By comparing the number of phosphorylation sites and spectrum counts of WT and *snf1Δ* (Figure S5), we performed mutagenesis and deletion of the potential phosphorylation sites and regions in four proteasome subunits, i.e., Rpn8, Pre10, Rpn13, and Rpn11 (Table S2). None of the mutated phosphorylation sites caused defects in proteasome condensate dissolution. Therefore, an optimized strategy will need to tease out the direct effector proteins that regulate proteasome condensate dissolution and phosphorylation status in the AMPK pathway.

The AMPK pathway plays a major role in regulating cellular energy homeostasis. Dysfunction of AMPK has been observed in various diseases, such as cancer, metabolic disorders, and aging (JEON 2016). Therefore, AMPK has attracted widespread interest as a therapeutic target for disease treatment. Our results indicate that AMPK plays an important role in proteasome condensate dissolution by interacting with the proteasome RP and altering proteasome phosphorylation status and condensate solubility, which is likely relevant to proteasome condensate regulation by AMPK in humans. Moreover, repurposing AMPK-modulating drugs for proteasome regulation could offer new strategies to tackle drug resistance in proteasome malfunction-related disease treatments.

## Materials and Methods

### Yeast strains and cell cultures

Yeast strains used in this study are listed in Table S3. Yeast cells were grown overnight in synthetic complete (SC) medium (LI AND HOCHSTRASSER 2022) at 30°C with vigorous agitation; tryptophan and uracil single or double dropout media were used for plasmid selection. Cultures were back-diluted in fresh SC medium and grown to mid-exponential phase. Cells were harvested and rinsed once with sterile ultrapure water and then resuspended in SC medium containing 0.025% glucose. The cells grown in low glucose cells were incubated for one and four days at 30°C with vigorous agitation. For glucose recovery (BUTCHER *et al*. 2025), the low-glucose starved cells were pelleted and rinsed once with sterile ultrapure water and then resuspended in fresh SC medium containing 2% glucose. The cells were incubated in this medium for 15 min and subjected to different experiments as described in the figures. For fluorescence microscopy, glucose-recovered cells were fixed with 2% formaldehyde at room temperature (RT) for 5 min. The fixed cells were pelleted, washed once with 0.1M KPO_4_ pH 6.5, and resuspended in 0.1M KPO_4_ pH 7.5 before imaging.

### Plasmid construction

Plasmids used in this study are listed in Table S4. Genes of interest were amplified from yeast genomic DNA and using restriction enzyme digestions and T4 DNA ligase (NEB)-mediated ligation were cloned into the following plasmids: pRS316 (SIKORSKI AND HIETER 1989), p424GPD, and p426GPD (MUMBERG *et al*. 1995). The plasmids were verified by enzymatic digestion and DNA sequencing. QuikChange site-directed mutagenesis (Agilent) was performed to generate *snf1-T210A*, *snf1-T210D*, *snf1-K84R*, and *snf1-G53R* mutants.

### Fluorescence microscopy

Yeast cells were visualized on an Axioskop microscope (Carl Zeiss) equipped with a Plan-Apochromat 100×/1.40 NA oil DIC objective lens, a CCD camera (AxioCam MRm; Carl Zeiss), and an HBO100W/2 light source. Images were taken using AxioVision software with an auto-exposure setup. Yeast cells used for Figure S2 were visualized on a Nikon Ts2R inverted fluorescence microscope equipped with a CFI60 Plan Apochromat Lambda D 100x/1.45 NA oil DIC objective lens, a Nikon DS-Fi3 camera, and an LED light source. Images were taken using NIS-Elements D software with a 400 ms exposure setup. Images were taken under a single focal plane and processed using Adobe Photoshop CC software. Percentage of cells with proteasome condensates were quantified using ImageJ. ANOVA single factor significance analysis was performed using Excel.

### Protein extraction and Western blotting

Total yeast proteins were extracted using the post-alkaline extraction method (KUSHNIROV 2000), and Western blotting was performed as described previously with minor changes (LI *et al*. 2016). One optical density unit at 600 nm (OD_600_) of cells were harvested at 8,000 rpm for 1 min and rinsed once with sterile ultrapure water at room temperature (RT). For phosphorylation analyses, cells were incubated in 400 µl 0.1 M NaOH with PhosSTOP phosphatase inhibitor cocktail tablets (Millipore Sigma, catalog # 4906837001) for 5 min at RT and pelleted at 10,000 rpm for 2 min. The alkali-treated cells were resuspended in 100 µl SDS sample buffer (10% glycerol, 2% SDS, 0.1 M DTT, 62.5 mM Tris-HCl pH 6.8, 4% 2-mercaptoethanol, 0.008% Bromophenol Blue, PhosSTOP phosphatase inhibitor) and heated at 100°C for 5 min. Cell debris was removed by centrifugation at 15,000 rpm for 1 min. Equal volumes of the supernatants were loaded onto 10% (v/v) SDS-PAGE gels with and without adding Mn^2+^-Phos-tag to the resolving gels. Additionally, the Phos-tag SDS PAGE gels were incubated with transfer buffer (20% methanol, 0.3% trizma base, 1.44% glycine, 0.01% SDS) containing 10 mM EDTA and then incubated in fresh transfer buffer before the transfer of proteins to PVDF membranes (EMD Millipore, catalog # IPVH00010).

The membranes were incubated with JL-8 anti-GFP monoclonal antibody (Clontech catalog # 632381) at 1:2,000 dilution or an anti-Pgk1 monoclonal antibody (Thermo Fisher Scientific, catalog # 459250) at a 1:10,000 dilution. Primary antibody binding was followed by anti-mouse-IgG (Cytiva, catalog # NXA931) secondary antibody conjugated to horseradish peroxidase at a 1:10,000 dilution. The membranes were incubated in ECL detection reagent (MRUK AND CHENG 2011). The ECL signals were detected using film (Thomas Scientific, catalog #1141J52).

### GFP-Trap immunoprecipitation of GFP-tagged proteasomes

GFP-Trap immunopurifications were performed as described previously (LI AND HOCHSTRASSER 2022). Briefly, 35 OD_600_ units of cells expressing GFP-tagged proteasome subunits were harvested and rinsed once with ice-cold sterile ultrapure water. Cells were resuspended in lysis buffer (20 mM Tris pH 7.5, 0.5 mM EDTA pH 8, 200 mM NaCl, 10% glycerol, 1 mM PMSF, 10 mM NEM, Roche cOmplete mini protease inhibitor catalog # 11836153001). Total cell lysates were prepared by beating with acid-washed glass beads (Sigma-Aldrich, catalog # G8772). The cell membranes were solubilized by adding 0.1% Triton X-100 to the lysates and incubated on ice for 30 min. Crude cell lysates were cleared at 16,000 x *g* for 10 min at 4°C. Cleared cell lysates were incubated with GFP-Trap agarose resin slurry for 1 h. The resin was washed three times with the lysis buffer containing 0.02% Triton X-100, resuspended in 2xSDS sample buffer, and then incubated at 42°C for 10 min to elute the bound proteins. The elutes were analyzed by Western blotting as above with monoclonal anti-HA antibody (Sigma, catalog # H9658) at a 1:2,000 dilution as well as with JL-8 anti-GFP and anti-Pgk1 monoclonal antibodies.

### *In vivo* proteasome condensate solubility test

Yeast cell treatment with 1,6-hexanediol (1,6-HD) was performed as described previously (KROSCHWALD *et al*. 2017). Low glucose-grown yeast cells were harvested and treated with a final concentration of 6% 1,6-hexanediol (1,6-HD) and 10 µg/ml digitonin in the media on day one, and 10% 1,6-HD and 10 µg/ml digitonin on day four. The cells were incubated at RT for 20 min and fixed with 2% formaldehyde before imaging by fluorescence microscopy as described above.

### Proteasome affinity purification and gel staining

Proteasome affinity purification was performed using yeast cells expressing Rpn11-3xFlag as described previously (SULTANA *et al*. 2023). Briefly, chromosomally tagged *RPN11-3xFlag* cells were harvested following incubation at the indicated growth conditions and ground to fine powder with liquid nitrogen. Cell powders were resuspended in buffer A (50 mM Tris-HCl pH 7.5, 150 mM NaCl, 10% glycerol, 5 mM MgCl_2_, 5 mM ATP, Roche cOmplete EDTA-free protease inhibitor, catalog #1872580001, PhosSTOP phosphatase inhibitor), and incubated on ice for 15 min. Cell debris were pelleted at 30,000 x *g* for 20 min at 4°C. Total protein concentration was determined with a Pierce BCA protein assay kit (Thermo Scientific, catalog #23227). The supernatants were incubated with anti-FLAG M2 affinity gel (Sigma, catalog #A2220) for 2 h on a rotator at 4°C. The resin was washed twice with buffer A for 10 min and then incubated with 3 resin volumes of 200 µg/ml 3xFLAG peptide (Sigma, catalog #F4799) for 45 min to elute proteasome complexes. Proteasomes were concentrated with 100 K MWCO centrifugal filters (Merck Millipore, catalog #UFC510024) and quantified with a BSA standard using a G:Box Chemi HR16 imager (Syngene).

Proteasome staining with Pro-Q® Diamond phosphoprotein gel stain (Invitrogen, catalog # P33301) and SYPRO® Ruby protein gel stain (Invitrogen, catalog # S12001) were performed according to the manufacturer’s protocols. Briefly, 10 µg of proteasomes were resolved in 12% SDS-PAGE gels. The gels were fixed twice with a solution of 50% methanol and 10% acetic acid with gentle agitation at RT for 30 min per fixation. The fixation step was to ensure the removal of SDS from the gels. The fixed gels were washed three times with ultrapure water for 10 min per wash to remove methanol and acetic acid from the gels. Then the gels were stained with Pro-Q® Diamond phosphoprotein gel stain with gentle agitation in the dark for 1 h and destained three times with a solution of 20% acetonitrile and 50 mM sodium acetate (pH 4.0) with gentle agitation at RT in the dark for 30 min each. The gels were washed twice with ultrapure water at RT for 5 min per wash before imaging with a Typhoon biomolecular imager (Amersham Biosciences).

After phosphoprotein gel stain imaging, the gels were incubated with SYPRO® Ruby protein gel stain solution overnight at RT. The stained gels were washed with a solution of 10% methanol and 7% acetic acid for 30 min and rinsed twice with ultrapure water for 5 min each before imaging with the BioRad ChemiDoc MP Imaging System.

### Affinity purification of Snf1 complexes

Yeast cells expressing Snf1-3xFlag from the chromosomally tagged *SNF1* locus were used for Snf1 complex affinity purification. Cells were harvested at the indicated growth conditions and ground to a fine powder with liquid nitrogen. Cell powders were resuspended in modified IPP150 buffer (10 mM Tris-HCl, pH 8.0, 150 mM NaCl, 0.1% Nonidet P-40, 5% Glycerol, and Roche cOmplete EDTA-free protease inhibitor, catalog #1872580001) (NATH *et al*. 2002). The lysed cells were incubated on ice for 15 min and followed the same protocol for proteasome purification above. Snf1 complexes were concentrated with 10 K MWCO centrifugal filters (Merck Millipore, catalog # UFC501024) and quantified with a BSA standard using a G:Box Chemi HR16 imager (Syngene). The purified Snf1 complexes were resolved in 10% SDS-PAGE gels. The gel slices were fixed before liquid chromatography-tandem mass spectrometry (LC-MS/MS) analysis (see below).

### In-gel digestion

Gel bands containing the Snf1 complexes or proteasomes were cut into small pieces and rinsed with 1 ml water on a tilt-table for 10 min. The bands were then washed for 20 min with 1 ml 50% acetonitrile (ACN)/100 mM NH_4_HCO_3_ (ammonium bicarbonate, ABC). The samples were reduced by the addition of 4.5 mM dithiothreitol (DTT) (sufficient volume to cover gel pieces) in 100 mM ABC with incubation at 37°C for 20 minutes. The DTT solution was removed, and the samples were cooled to room temperature. Samples were alkylated by the addition of an equal volume of 10 mM iodoacetamide (IAN) in 100 mM ABC with incubation at room temperature in the dark for 20 minutes. The IAN solution was removed, and the gels were washed for 20 min with 1 ml 50% ACN/100 mM ABC, then washed for 20 min with 1 ml 50% ACN/25 mM ABC. The gels were briefly dried in a SpeedVac, then resuspended in 100-150 µl of 25 mM ABC containing 200-300 ng of sequencing grade trypsin (Promega, V5111) (volume sufficient to cover gels) and incubated at 37°C for 16 hours. The samples that were to be submitted for phosphopeptide enrichment had more protein and so were digested with 500 ng of trypsin. The supernatant containing the released tryptic peptides were moved to a new Eppendorf tube. Residual peptides remaining in the gel were extracted by the addition of approximately 3 volumes of 80% ACN/0.1% trifluoroacetic acid and combined with the previous supernatant, and the peptides were dried in a SpeedVac. For unenriched samples, peptides were dissolved in 23 µl MS loading buffer (2% ACN, 0.2% TFA in water), with 5 µl injected for LC-MS/MS analysis. For select samples, after extraction, phosphorylated peptides were enriched using TiO2 (titanium dioxide) TopTips (TT1TIO.96; Glygen, Columbia, MD) with a slightly modified manufacturer protocol. Briefly, the manufacturer protocol was utilized with the addition of 70 mM L-glutamic acid in the loading buffer (65% acetonitrile, 2% trifluoroacetic acid). Bound phosphopeptides on the TiO2 resin were washed with 65% acetonitrile, 2% trifluoroacetic acid, and eluted with 2% ammonium hydroxide solution in water at pH 12. The enriched fraction was acidified, and both enriched (EN) and flowthrough (FT) fractions were dried by SpeedVac prior to dissolving in MS loading buffer for LC–MS/MS analysis.

### LC-MS/MS on the Thermo Scientific Q Exactive Plus

LC-MS/MS analysis was performed as previously described (LI AND HOCHSTRASSER 2022) at the Keck Mass Spectrometry & Proteomics Resource at the Yale School of Medicine on a Thermo Scientific Q Exactive Plus equipped with a Waters nanoAcquity UPLC system utilizing a binary solvent system (A: 100% water, 0.1% formic acid; B: 100% acetonitrile, 0.1% formic acid). Trapping was performed at 5µl/min, 99.5% Buffer A for 3 min using a Waters ACQUITY UPLC M-Class Symmetry C18 Trap Column (100Å, 5 µm, 180 µm x 20 mm, 2G, V/M). Peptides were separated at 37°C using a Waters ACQUITY UPLC M-Class Peptide BEH C18 Column (130Å, 1.7 µm, 75 µm X 250 mm) and eluted at 300 nl/min with the following gradient: 3% buffer B at initial conditions; 5% B at 1 minute; 25% B at 90 minutes; 50% B at 110 minutes; 90% B at 115 minutes; 90% B at 120 min; return to initial conditions at 125 minutes. Full MS spectra were acquired in profile mode over the 300-1,700 m/z range using 1 microscan, 70,000 resolution, AGC target of 3E6, and a maximum injection time of 45 ms. Data dependent MS/MS scans were acquired in centroid mode on the top 20 precursors per MS scan using 1 microscan, 17,500 resolution, AGC target of 1E5, maximum injection time of 100 ms, and an isolation window of 1.7 m/z. Precursors were fragmented by HCD activation with a collision energy of 28%. MS/MS spectra were collected on species with an intensity threshold of 1E4, charge states 2-6, and peptide match preferred. Dynamic exclusion was set to 20 seconds.

### Peptide identification

Tandem mass spectra were extracted by Proteome Discoverer software (version 2.2.0.388, Thermo Scientific) and searched in-house using the Mascot algorithm (version 2.7.0, Matrix Science). The data were searched against the SWISS-PROT database with taxonomy restricted to *Saccharomyces cerevisiae* (v20211116; 7,921 sequences). Search parameters included trypsin digestion with up to 2 missed cleavages, peptide mass tolerance of 10 ppm, and MS/MS fragment tolerance of 0.02 Da. Cysteine carbamidomethylation was configured as a fixed modification. Methionine oxidation; phosphorylation of serine, threonine and tyrosine; propionamide adduct to cysteine; and GG dipeptide (ubiquitin residue) on lysine were configured as variable modifications. Normal and decoy database searches were run, with the confidence level was set to 95% (p<0.05). Scaffold Q+S (version 5.1.2, Proteome Software Inc.) was used to validate MS/MS based peptide and protein identifications. Peptide identifications were accepted if they could be established at greater than 95.0% probability by the Scaffold Local FDR algorithm. Protein identifications were accepted if they could be established at greater than 99.0% probability and contained at least 2 identified peptides. The phosphopeptide enriched samples were further analyzed with Scaffold PTM (version 4.0.2, Proteome Software Inc.).

## Data availability

The proteomic data from this study has been deposited in Integrated Proteome Resources with ID number IPX0012452000. A full list of proteins identified in the AMPK complex (Figure S4) and proteasomes (Figure S6) by mass spectrometry analysis is provided in Table S5. All other data are included in the manuscript.

## Acknowledgements

We thank the Keck MS & Proteomics Resource at the Yale school of Medicine for providing the necessary mass spectrometers and the accompanying biotechnology tools funded in part by the YSM and NIH (S10OD02365101A1, S10OD019967, and S10OD018034).

## Study Funding

The work was supported by the startup fund and provost seed grant from Florida Institute of Technology to JL and NIH grant GM136325 to MH.

## Conflicts of interest

The author(s) declare no conflict of interest.

## Author contributions

JL: conceptualization, methodology, investigation, data curation, supervision, funding acquisition, project administration, writing-original draft, and writing-review and editing; CB and KV: investigation; MH: conceptualization, supervision, funding acquisition, project administration, and writing-review and editing.

